# Matrix alignment and density modulate YAP-mediated T-cell immune suppression

**DOI:** 10.1101/2024.03.19.585707

**Authors:** Jiranuwat Sapudom, Aseel Alatoom, Paul Tipay, Jeremy CM Teo

## Abstract

T-cells navigate through various mechanical environments within the body, adapting their behavior in response to these cues. An altered extracellular matrix (ECM) characterized by increased density and enhanced fibril alignment, as observed in cancer tissues, can significantly impact essential T-cell functions critical for immune responses. In this study, we used 3D collagen matrices with controlled density and fibril alignment to investigate T-cell migration, activation, and proliferation. Our results revealed that dense and aligned collagen matrices suppress T-cell activation through enhanced YAP signaling. By inhibiting YAP signaling, we demonstrated that T-cell activation within these challenging microenvironments improved, suggesting potential strategies to enhance the efficacy of immunotherapy by modulating T-cell responses in dense and aligned ECMs. Overall, our study deepens our understanding of T-cell mechanobiology within 3D relevant cellular microenvironments and provides insights into countering ECM-induced T-cell immunosuppression in diseases such as cancer.

**Graphical abstract:** Dense and aligned extracellular matrices suppress T-cell activation via YAP signaling, affecting immunotherapy efficacy in diseases such as cancer.

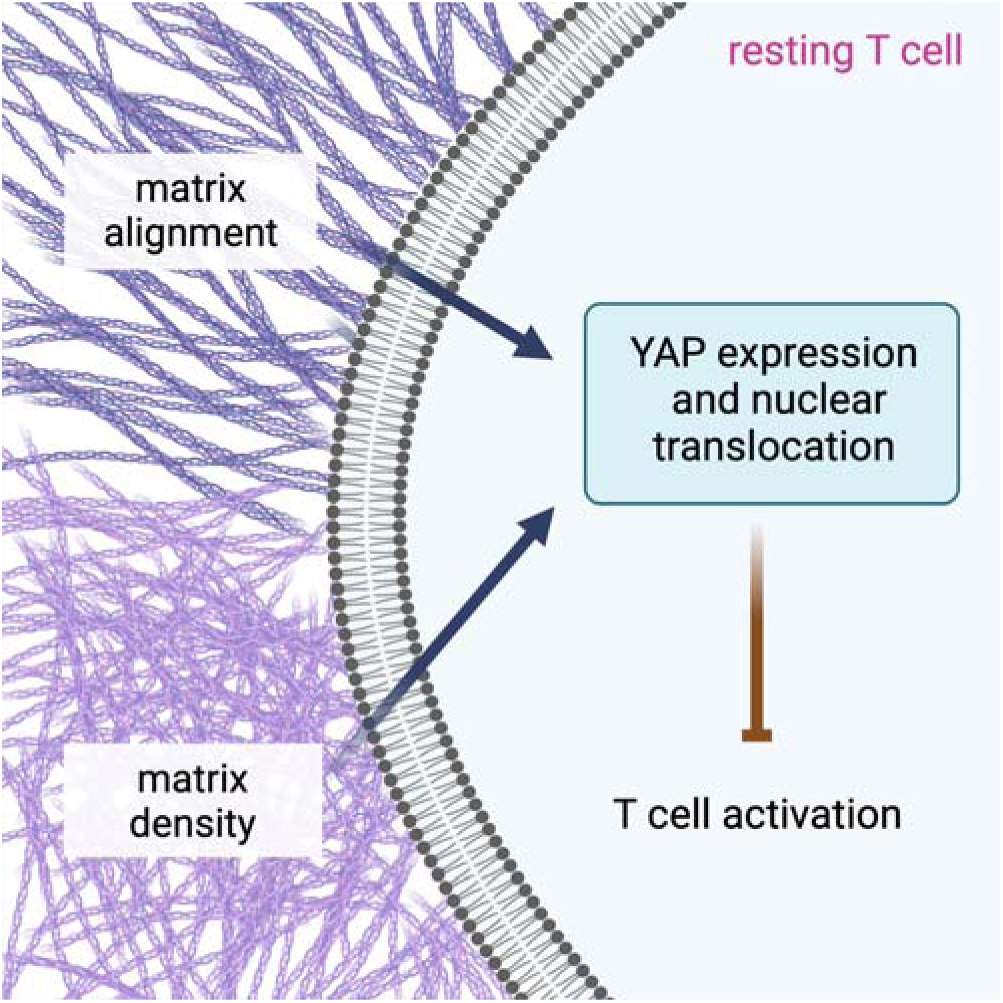

## 1. Introduction

As T-cells navigate through the body, they traverse a myriad of mechanical microenvironments, seamlessly transitioning from the bloodstream through the intricate network of blood vessels and into the complex, fibrillar structures of tissues. This journey is likely underpinned by the principle of cellular mechanotransduction [1–3], a process that enables T-cells to fine-tune their behavior, optimizing immunological responses and operational efficacy. Such adaptability is underscored by observations that T-cells alter their migration patterns in response to the diverse mechanical cues encountered across different tissue landscapes [4–6]. Although the role of cytokines in modulating immune responses has been extensively documented, the influence of the biophysical properties of tissues on T-cell functionality has not been fully explored; however, a comprehensive understanding of T-cell behavior, particularly under disease conditions, is imperative. This becomes even more critical in the context of pathological states, where alterations in the extracellular matrix (ECM), such as those observed in tumors and scar formations or as part of the aging process, are pronounced and impactful [7–9].

Tumors are cancer cells that manipulate a variety of stromal cells to promote their growth and spread [10]. These malignant cells compel adjacent stromal cells to form what is termed the tumor microenvironment (TME), which is characterized by remodeling of the extracellular matrix (ECM) and highlighted by the presence of aligned and dense fibrillar collagen structures. The TME aids in the migration of cancerous cells and assists in evading immune detection by impeding T-cell infiltration [11–13]. Histological examinations of solid tumors revealed that T-cells are scarce and predominantly found in the adjacent tumor stroma [14–16]. Additionally, the obstruction of T-cell infiltration presents a considerable obstacle for T-cell-based immunotherapies [17, 18]. Elucidating the processes that bolster immunosuppression within tumors could significantly improve these therapeutic approaches.

Recent studies on the mechanobiology of T-cells have employed hydrogel systems, including polyacrylamide, polydimethylsiloxane, and alginate, which immobilize ligands on their surfaces with tunable stiffness. These systems are designed to mimic cell□cell interactions between antigen-presenting cells (APCs) and T-cells, advancing our understanding of the mechanical cues central to APC and T-cell interactions [19–21]. Despite these advancements, the effects of tissue alterations during cancer progression on T-cell activation and function remain poorly understood. Although recent efforts have aimed to replicate the stiffness of cancerous tissue using alginate hydrogels [22], such hydrogels do not possess the fibrillar features inherent in native tissues.

Employing 3D fibrillar collagen matrices offers significant advantages for biomimicking tissue structures, enhancing our understanding of cellular interactions within a surrogate tissue context [23, 24]. These matrices can be customized in terms of fibril microstructure, crosslinking density, and integration with other ECM molecules [25], enabling the simulation of both homeostatic and pathophysiological conditions. Recent initiatives have utilized 3D collagen matrices with aligned collagen fibrils to study T-cell migration [13, 26], adjustable collagen density to examine T-cell activity [11], and tunable viscoelasticity to modulate T-cell subtypes [27]. However, a comprehensive study to elucidate T-cell activation and functions, alongside a mechanistic exploration of T-cell mechanosensing in response to matrix alterations, is still needed.

In this study, we aimed to replicate the 3D fibrillar configurations by adjusting the matrix density and fibrillar alignment. By utilizing these reconstituted matrices with varying microarchitectures, we investigated T-cell activation and function, including migration and proliferation. To gain mechanistic insights into the suppression of T-cell activation, we inhibited various biological pathways implicated in cell mechanosensing and studied the immune response. Our work provides a deeper understanding of how T-cell activation and function are affected by aberrant changes in the ECM fibrillar microstructure.

## 2. Experimental Section

### 2.1. Reconstitution of 3D collagen matrix

Three-dimensional (3D) collagen matrices were prepared following methods described in previous publications [28]. Briefly, rat tail type I collagen (Advanced BioMatrix, Carlsbad, CA, USA) was combined with 500 mM phosphate buffer (Sigma-Aldrich, Darmstadt, Germany) and 0.1% acetic acid (Sigma-Aldrich, Darmstadt, Germany) to obtain final collagen concentrations of 1 mg/ml and 3 mg/ml, respectively, to replicate loose and dense matrices. The prepared collagen solution was then transferred onto a glutaraldehyde-coated coverslip [24, 29–31]. Collagen fibrillation was induced at 37°C under 5% CO□ and 95% humidity for 30 minutes. To reconstitute 3D collagen matrices with aligned fibrils, coverslips containing the collagen solution were positioned on a custom 3D-printed stage with a 30° incline [32], and collagen fibrillation was initiated under the same conditions.

### 2.2. Topological and mechanical characterization of reconstituted 3D collagen matrices

Before initiating cell culture, the topological and mechanical properties of the cell-free collagen matrices were evaluated. For topological assessments, the 3D collagen matrices were stained with 50 µM 5-(6)-carboxytetramethylrhodamine succinimidyl ester (TAMRA-SE; Sigma-Aldrich, Darmstadt, Germany) overnight at room temperature and subsequently rinsed with phosphate-buffered saline (PBS; Sigma-Aldrich, Darmstadt, Germany). The matrices were then imaged using a confocal laser scanning microscope (Leica STED SP8 microscope; Leica Microsystems, Wetzlar, Germany) equipped with a 63× oil-immersion objective (NA 1.3). Stacked images were acquired at four random positions per matrix, with an image resolution of 1024 × 1024 pixel (xyz-voxel size: 0.13 × 0.13 × 10 µm). Pore size and fibril diameter were quantified utilizing a custom image analysis toolbox as described in a previous study [29]. Additionally, the fibril alignment distribution and coherence index (alignment index) were analyzed with OrientationJ [33], an ImageJ (NIH, Bethesda, Maryland, USA) plugin. The quantification was performed at four randomly selected positions of each sample from four independent samples.

In addition to topological analyses, mechanical characterization was performed using ElastoSens™ Bio 2 (Rheolution Inc., Montréal, Canada) through a nondestructive rheological approach, as detailed in prior publications [32]. These evaluations of topological and mechanical properties were conducted in four replicates.

### 2.3. Cell Culture

Human Jurkat T-cells (E6-1 clone; ATCC, Manassas, VA, USA) were cultured in RPMI-1640 medium supplemented with 10% fetal bovine serum (FBS), 1% penicillin/streptomycin, sodium pyruvate, HEPES, and 0.01% beta-mercaptoethanol. The cells were incubated under standard culture conditions at 37°C with 5% CO□ and 95% humidity. All cell culture reagents were obtained from Gibco (Invitrogen, Waltham, Massachusetts, USA).

### 2.4. Activation of T-cells

A total of 1×10^5^ Jurkat T-cells were seeded onto 3D collagen matrices and incubated overnight to facilitate their infiltration into the matrices. Subsequently, the cells were activated by incubation in RPMI-1640 medium supplemented with 20 ng/mL phorbol 12-myristate 13-acetate (PMA) and 1 µg/mL ionomycin under standard culture conditions as described previously [34]. All activators were procured from Sigma-Aldrich, Darmstadt, Germany. After 3 hours of incubation, the activation medium was discarded, and the cells were washed three times with PBS (Sigma-Aldrich, Darmstadt, Germany). Then, the cells were maintained in RPMI-1640 medium without the activators for 3 days before subsequent analyses were performed.

### 2.5. Visualization of cells and quantification of the actin cytoskeleton

To visualize the morphology of resting and activated T-cells within 3D collagen matrices, cells were stained with 1 µM carboxyfluorescein succinimidyl ester (CFSE) dye (BioLegend, San Diego, CA, USA) for 15 minutes under standard cell culture conditions, followed by fixation with 4% paraformaldehyde (PFA) for 10 minutes. Imaging was performed using a confocal laser scanning microscope (Leica STED SP8; Leica Microsystems, Wetzlar, Germany) equipped with a 63x oil-immersion objective.

For actin cytoskeleton staining, after fixation with 4% paraformaldehyde for 10 minutes, the cells were permeabilized with 0.1% Triton X-100 for 10 minutes at room temperature. Subsequently, they were stained overnight at 4°C with phalloidin conjugated to Alexa Fluor 488 (1:250 dilution in PBS; Invitrogen, Waltham, Massachusetts, USA) and DAPI (1:10,000 dilution in PBS). These cells were then imaged using the same confocal microscope with a 63× oil-immersion objective. To quantify actin, both resting and activated cells were retrieved from the 3D collagen matrices using 4 mg/mL collagenase Type VI, followed by fixation, permeabilization, and staining with phalloidin conjugated to Alexa Fluor 488. The actin content was quantified using an Attune NxT Flow Cytometer (ThermoFisher Scientific, Waltham, Massachusetts, USA), and the geometric mean fluorescence intensity (gMFI) was calculated using FlowJo software (version 10.10; BD Bioscience, NJ, USA). The experiments were conducted in four replicates.

### 2.6. 3D live-cell imaging and quantitative label-free single-cell tracking

3D live cell imaging was conducted using a bright-field microscope (DMi8 S microscope platform; Leica Microsystems, Wetzlar, Germany) equipped with an incubation chamber and motorized stage. The incubation chamber was maintained under conditions analogous to those of standard cell culture environments: 37°C, 95% humidity, and 5% CO_2_. Sequential stacked images were captured using a 10× objective (Leica Microsystems, Wetzlar, Germany) in bright-field mode, with image acquisition every 7 µm in the z-direction across 65 z-layers, totaling a 455 µm z-depth. This process occurred at 6-minute intervals over a 24-hour period. The resolution of the images was 1388 × 1040 pixels, with x- and y-voxel sizes of 0.87 µm.

Cell migration speed and directionality were evaluated using custom-developed automated 3D label-free cell tracking software [35]. The average cell migration speed and directionality were calculated for at least 73 cells under each matrix condition for both resting and activated T-cells. Migration directionality was quantified by comparing the Euclidean and accumulated distances of the cell trajectory, where a directionality of 0 indicates random migration and a directionality of 1 signifies perfectly linear migration from the starting to the endpoint.

### 2.7. Quantitative analysis of T-cell activation through the expression of cell surface markers and intracellular cytokine production

For the quantification of cell activation, cells were harvested from their matrices using liberase DL research grade (24 U; Roche, Basel, Switzerland) and stained with the following antibodies (diluted 1:500 in PBS) for 30 min on ice: mouse-anti-human CD25 antibody conjugated with brilliant violet 605^TM^ (clone: BC96), mouse-anti-human CD44 antibody conjugated with FITC (clone: BJ18), mouse-anti-human CD69 antibody conjugated with PE/Fire^TM^640 (clone: FN50), and mouse-anti-human PD1 antibody conjugated with brilliant violet 421^TM^ (clone: EH12.2H7), all of which were obtained from BioLegend, San Diego, CA, USA. The expression of cell surface markers was quantified by measuring the gMFI using an Attune NxT Flow Cytometer equipped with an autosampler (ThermoFisher Scientific, Waltham, Massachusetts, USA). The experiment was conducted with four independent replicates.

T-cells were cultured in RPMI-1640 media supplemented with 5 µg/mL brefeldin A (BioLegend, San Diego, CA, USA). After 24 hours of cultivation, the cells were harvested from the matrices by digesting the collagen with liberase DL research grade (24 U; Roche, Basel, Switzerland). The cells were then fixed with 4% PFA, permeabilized with 0.1% Triton X-100, and blocked with 1% bovine serum albumin (BSA). The sections were subsequently stained with a mouse anti-human IFNγ antibody conjugated with Brilliant Violet 605^TM^ (dilution: 1:500; clone: 4S. B3), and rat anti-human IL-2 antibody conjugated with APC/Cyanine7 (dilution: 1:500; clone: MQ1-17H12) for 2 hours at room temperature. After the cells were washed with PBS, the fluorescence intensity of the stained IFNγ and IL-2 antibodies was quantified using an Attune NxT Flow Cytometer equipped with an autosampler (ThermoFisher Scientific, Waltham, Massachusetts, USA). The experiment was conducted with four independent replicates.

### 2.8. Quantitative analysis of cytokine secretion

After 3 days of culture of activated T-cells under various matrix conditions, the cell culture supernatants were collected for cytokine analysis. The secretion profiles of IL-2, TNF-α, and IFNγ were quantified using a bead-based multiplex immunoassay according to the manufacturer’s protocols. The samples were then analyzed with an Attune NxT Flow Cytometer (ThermoFisher Scientific, Waltham, Massachusetts, USA). Data analysis was conducted using LEGENDplex™ data analysis software (BioLegend, San Diego, CA, USA). The experiments were carried out in four independent replicates.

### 2.9. Quantitative analysis of T-cell proliferation

Before seeding T-cells onto 3D collagen matrices and subsequent activation, the cells were stained with 2.5 µM Violet Tag-It dye (BioLegend, San Diego, CA, USA) for 15 minutes at 37°C, 95% humidity, and 5% CO_2_. Subsequently, the cells were activated using PMA and ionomycin as described previously in Section 2.4. For the quantitative analysis of T-cell proliferation, Violet Tag-it-positive cells were analyzed for their fluorescence intensity using an Attune NxT Flow Cytometer equipped with an autosampler (ThermoFisher Scientific, Waltham, Massachusetts, USA). The expansion index (EI) was calculated using the cell proliferation module in FlowJo software (version 10.10; BD Bioscience, NJ, USA). The experiment was conducted in four independent replicates.

### 2.10. Inhibition of mechanotransduction-related pathways

Prior to chemically activating T-cells with PMA and ionomycin, T-cells were treated with various mechanotransduction-associated inhibitors, namely, cytochalasin B (1 µM), blebbistatin (5 µM), Y-27632 (1 µM), colchicine (1 µM), verteporfin (1 µM), GsMTx4 (5 µM), an anti-integrin β1 antibody (2 µg/mL), manganese chloride (0.5 mM MnCl_2_), and a PI3K inhibitor (1 µM), in RPMI-1640 cell culture media. All inhibitors were purchased from TOCRIS, Minneapolis, MN, USA. After three hours, the inhibitors were removed, and the cells were washed with PBS prior to activation, as detailed in Section 2.4. Subsequently, the cells were analyzed for activation markers and intracellular cytokine production following the protocols described in Section 2.7. The experiment was conducted in four independent replicates.

### 2.11. YAP staining and quantification

Resting T-cells were cultured in RPMI-1640 media supplemented with 5 µg/mL brefeldin A (BioLegend, San Diego, CA, USA). After 24 hours of cultivation, the cells were harvested from the matrices by digesting the collagen with liberase DL research grade (24 U; Roche, Basel, Switzerland). The cells were then fixed with 4% PFA, permeabilized with 0.1% Triton X-100, and blocked with 1% bovine serum albumin (BSA). The sections were subsequently stained with a rat anti-human YAP antibody conjugated with PE (dilution: 1:500; clone W19260A; Biolegend, San Diego, CA, USA) for 2 hours at room temperature. After washing, YAP expression was quantified by measuring the fluorescence intensity using an Attune NxT Flow Cytometer equipped with an autosampler (ThermoFisher Scientific, Waltham, Massachusetts, USA). The experiment was conducted with four independent replicates.

To quantify YAP nuclear translocation, resting T-cells were cultured as described above and subsequently processed in a similar manner. For staining, different dilutions of the YAP antibody conjugated with PE (dilution: 1:500; clone W19260A) and DAPI (dilution: 1:10,000; Invitrogen, Waltham, Massachusetts, USA) were applied for 2 hours at room temperature. Imaging was conducted using a Lionheart FX automated microscope (BioTek, Winooski, VT, USA) equipped with a 10× objective lens. YAP nuclear translocation was assessed using the built-in analysis software Lionheart FX. This analysis segmented the cytoplasm and nucleus based on the YAP fluorescence signal and the DAPI signal, respectively. The fluorescence intensity of YAP in the segmented cytoplasm and nucleus was quantified as a ratio. A minimum of 109 cells were analyzed for each experimental condition.

### 2.12. Statistical analysis

The experiments were conducted in at least four replicates, unless otherwise specified. The data are presented as the mean, and the error bars represent the standard deviation (SD). Levels of statistical significance were assessed using one-way ANOVA followed by Dunn’s test with GraphPad Prism 8 (GraphPad Software, San Diego, CA, USA). The levels of significance were established as follows: *p < 0.05, **p < 0.01, ***p < 0.001, ****p < 0.0001.

## 3. Results and Discussion

An increase in collagen density and improved alignment of fibrils are hallmark characteristics observed in the pathological landscapes of fibrotic tissues within cancer microenvironments, scars, and aged tissues [7–9]. Within these environments, T-cells play a crucial role in maintaining homeostasis and modulating pathological processes. To effectively execute their functions, these cells must adapt to the varying microstructural and compositional nuances of the surrogate extracellular matrix (ECM) in different tissues.

Given the limitations of the current tissue engineering approaches in replicating 3D collagen matrices with fiber alignment akin to those found in pathological settings, we utilized our straightforward, high-throughput technique using 3D-printed inclined surfaces, as previously reported [32]. Subsequently, we investigated the mechanobiological responses of T-cells to these engineered matrices, focusing on aspects such as activation status, migration, proliferation, surface marker expression, and cytokine secretion. In addition, we attempted to identify possible biological pathways that might be responsible for the suppression of T-cell activation, potentially allowing us to improve T-cell responses in the cancer microenvironment.

### 3.1. Reconstitution of loose and dense 3D collagen matrices with random and aligned fibrils

To reconstitute loose and dense matrices, collagen solutions were prepared at concentrations of 1 mg/ml and 3 mg/ml, respectively. Afterward, the collagen solution was transferred onto coated coverslips to initiate collagen fibrillation. To construct matrices with random collagen fibrils, coverslips were placed on planar surfaces. In contrast, matrices with fibril alignment were reconstituted by placing the coverslips onto 3D-printed inclined surfaces at a 30° angle, as previously reported [32].

As shown in **Figure 1A**, representative images depict loose and dense matrices with random and aligned collagen fibrils. Increasing the collagen concentration results in denser collagen matrices. Furthermore, pronounced collagen fibril alignment was observed at both concentrations. To validate these visual observations, we quantitatively analyzed the obtained images to gain insights into the distribution of collagen fibrils and their degree of alignment. The distribution of collagen fibril orientation revealed that collagen matrices fibrillated on inclined surfaces displayed a Gaussian distribution. In contrast, collagen matrices fibrillated on planar surfaces exhibited uniformly distributed fibril orientations, indicating that all fibril angles are likely randomly oriented (**Figure 1B**). We further quantified the fiber alignment index by calculating the coherence index, which ranges from 0 to 1, where 0 indicates a perfectly random distribution and 1 indicates perfect alignment. As shown in **Figure 1C**, the coherence indices were 0.21 ± 0.05 and 0.21 ± 0.03 for random collagen from the loose and dense matrices, respectively. This index increased to 0.60 ± 0.03 and 0.53 ± 0.08 for the aligned matrices in loose and dense conditions, respectively. These coherence indices for the matrices are in a similar range to those reported for human normal tissue (CI: 0.21 ± 0.10) [36] and cancer-treated extracellular matrix (ECM) (CI: 0.60 ± 0.04) [37].

**Figure 1:**
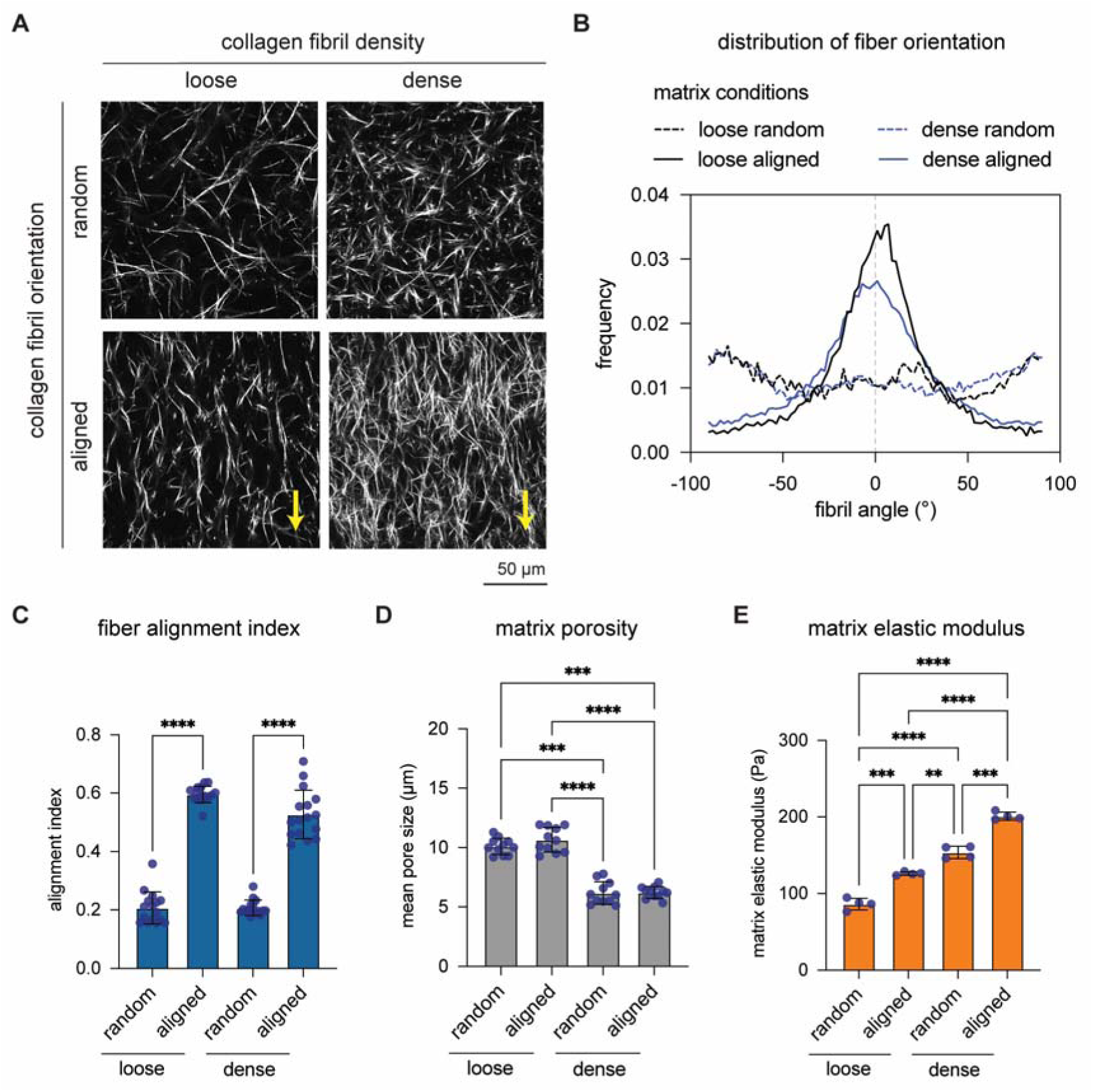
Characterization of reconstituted 3D collagen matrices. **(A)** Representative images of random and aligned fibrils within loose and dense 3D collagen matrices are presented. The arrow indicates the direction of collagen fibril alignment. (Scale bar = 50 µm). The topological parameters of these 3D collagen matrices were assessed using an image analysis toolbox, which included **(B)** the distribution of fibril orientation (0° is the direction of collagen fibril alignment), **(C)** the fibril alignment index, and **(D)** matrix porosity. The topological analysis was performed at four randomly selected positions of each sample from four independent samples. **(E)** The elastic modulus of the 3D collagen matrices was determined using a nondestructive rheological method in four independent replicates. The data are presented as the means ± SDs. Levels of statistical significance were assessed using one-way ANOVA followed by Dunn’s test. The levels of significance were established as follows: *p < 0.05, **p < 0.01, ***p < 0.001, ****p < 0.0001.

We also extended our analysis to investigate matrix porosity using a custom-made image analysis toolbox [29]. As depicted in **Figure 1D**, the pore size remained constant at diameters of 10.10 ± 0.68 µm and 10.66 ± 1.04 µm for loose collagen in random and aligned matrices, respectively. The pore size for dense collagen decreased to diameters of 6.17 ± 0.95 µm and 6.23 ± 0.50 µm, respectively. Thus, the pore size remained consistent regardless of the orientation of the collagen fibers. The bulk matrix elastic modulus was measured using a nondestructive rheometer, as illustrated in **Figure 1E**. The elastic modulus increased from 86.3 ± 7.6 Pa to 126.2 ± 2.23 Pa for loose collagen in random and aligned matrices, respectively, and from 153.8 ± 8.1 Pa to 200.9 ± 5.2 Pa for dense collagen, respectively. These data corroborate previous reports showing that an increase in matrix density enhances the matrix elastic modulus [31, 38]. Furthermore, the results indicate that matrix alignment significantly increases the matrix elastic modulus while maintaining similar porosity levels, as previously published [32, 39].

Overall, we reconstituted matrices with loose and dense 3D fibrillar collagen matrices featuring randomly distributed and aligned collagen fibrils. These well-defined matrices serve as valuable biomimetic models for investigating T-cell activation and function.

### 3.2. Activated T-cells exhibited increased protrusions and reduced actin content in aligned matrices

To investigate T-cell activation and function in various matrices, we used the Jurkat T-cell line, a widely recognized model for studying the T-cell mechanoresponse. This human lymphocytic cell line, Jurkat, was used to circumvent the biological variability and activation discrepancies inherent in primary cells. A recent report suggested that physical T-cell-APC interactions are less frequent in 3D biomimetic models, resulting in some T-cells remaining inactivated [31]. Therefore, we chemically activated T-cells using PMA and ionomycin, the latter of which is a calcium ionophore, to achieve T-cell activation. In conjunction with PMA, ionomycin significantly augments the activation of protein kinase C (PKC) [40]. This finding suggested that the role of ionomycin extends beyond the presumed activation of Ca^+2^/calmodulin-dependent signaling pathways. It also appears to induce phosphoinositide hydrolysis and consequent PKC activation in human T-cells [40]. This method of chemical activation is widely used in many studies [34, 41, 42] and bypasses T-cell receptor (TCR) activation, preventing its antigen specificity. The activation states of chemically activated T-cells and those activated through TCR signaling are similar, as demonstrated by transcriptional analysis [43, 44]. Furthermore, studies have shown that chemical activation using PMA and ionomycin can fully activate T-cells, as confirmed by sustained levels of IFN-γ, IL-2, and IL-4, which are cytokines commonly associated with T-cell activation [40, 41, 45]. By chemically activating genetically similar T-cells with PMA and ionomycin, we uniformly activated T-cells, allowing us to observe and confidently measure the impact of an altered matrix microenvironment on T-cell activation.

To observe morphological changes upon chemical activation, we visualized both resting and activated T-cells using a confocal microscope. As shown in **Figure 2A**, resting T-cells exhibited a smoother surface with very few protrusions under all matrix conditions, while the activated T-cells appeared larger with a rougher surface and exhibited pronounced protrusions. These protrusions have been identified as microvilli and invadosome-like structures in activated T-cells. They play roles in antigen probing and receptor triggering processes [46–50]. Moreover, the morphological appearance of activated T-cells suggested that they were activated in 3D microenvironments, and this effect appeared to be independent of the matrix conditions.

**Figure 2:**
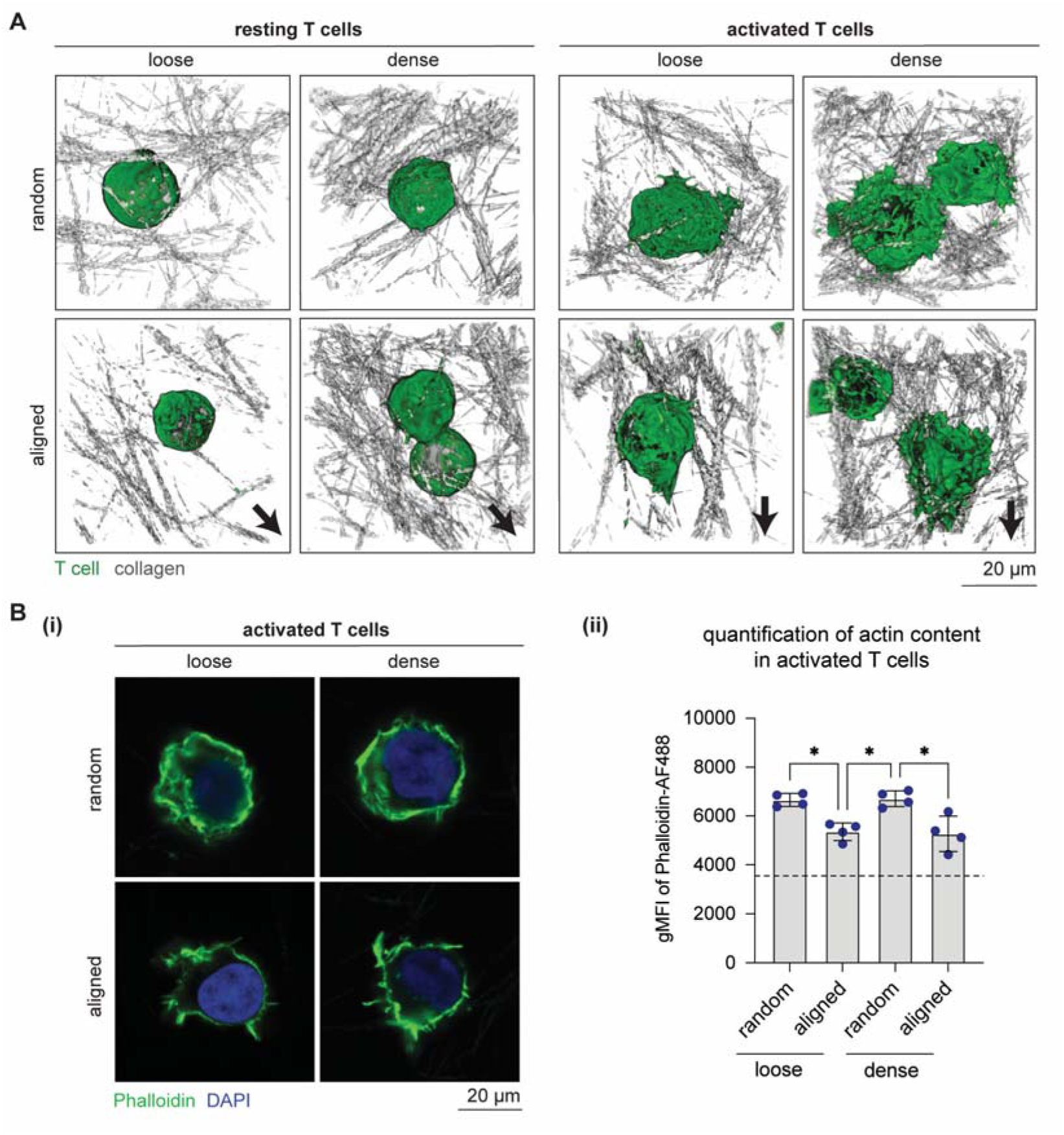
Morphology and actin content of T-cells under different matrix conditions. **(A)** Representative images depicting the morphology of T-cells within different 3D collagen matrices, rendered through CFSE staining (green) for T-cells and reflection imaging of collagen fibrils (gray). The arrow indicates the direction of collagen fibril alignment. (Scale bar = 20 µm) **(B(i))** Representative images of actin staining in activated T-cells using phalloidin conjugated to Alexa Fluor 488 (green) and DAPI staining for nuclei (blue). Quantitative analysis of actin content in activated T-cells using flow cytometry to measure the geometric mean fluorescent intensity (gMFI) of phalloidin-Alexa Fluor 488-stained cells. The data are presented as the means ± SDs. The dashed line indicates the average gMFI for resting cells across all conditions. Statistical significance was determined using one-way ANOVA with Dunn’s post hoc test, denoted as follows: *p < 0.05, **p < 0.01, ***p < 0.001, ****p < 0.0001. The experiments were conducted with four independent replicates.

As rapid F-actin polymerization is one of the hallmark indicators of T-cell activation [45, 51], we further investigated whether alterations in the matrix would affect the actin content of resting and activated T-cells. To investigate this possibility, we first visualized actin in activated T-cells by staining with phalloidin and imaging using a confocal microscope. As demonstrated in **Figure 2B(i)**, filamentous actin (F-actin) within the activated cells predominantly assembled in the cortex, with reduced quantities observed in the aligned matrices. Moreover, the change in cytosolic actin content was even less pronounced within activated T-cells in the aligned matrices. We further quantified these observations using flow cytometry analysis (**Figure 2B(ii)**), which revealed a significant reduction in actin content in cells within aligned matrices under both loose and dense conditions. Importantly, the actin content remained unchanged and decreased in resting T-cells, irrespective of the matrix conditions **(Supplementary Figure S1)**. These findings are consistent with previous studies that demonstrated decreased cell protrusions in aligned collagen matrices [13].

### 3.3. Matrix alignment reduces T-cell migratory capacity

The migration of T-cells is pivotal for scanning antigens presented by antigen-presenting cells and engaging in interactions with other cells crucial to the immune response. Thus, T-cell migration, characterized by nonproteolytic amoeboid movement, involves adaptive morphology, transient contact with surrogate ECM components, and navigation through matrix pores and plays an essential role in T-cell responses [52]. Increased density in breast cancer samples has been demonstrated to impede T-cell infiltration [11]. Similarly, studies have shown T-cell accumulation in regions of low collagen density within ex vivo tumor samples, with limited infiltration into collagen-dense regions [18, 53].

To investigate T-cell migration within 3D collagen matrices, live-cell imaging was conducted using a microscope equipped with a z-motor stage in bright-field mode, ensuring minimal cellular manipulation. Stacked images were collected at 6-minute intervals over 24 hours. For an in-depth analysis of T-cell migration characteristics, images were processed using custom-built quantitative label-free 3D single-cell tracking software [35]. Representative cell migration trajectories of resting and activated T-cells under different matrix conditions are illustrated in **Figure 3A**. Visual observation revealed that resting T-cells migrate faster and more randomly, whereas activated T-cells exhibit reduced migratory capacity. Specifically, the migration rate of activated T-cells within dense collagen matrices, whether with random or aligned fibrils, is notably decreased. We hypothesize that this observation reflects the immunological requirement wherein resting T-cells rapidly migrate to survey antigen-presenting cells (APCs) in peripheral tissues. Activated T-cells subsequently reduce their migration to remain in inflamed tissues, which facilitates their proper activation and subsequent differentiation into effector or memory T-cell subsets [4–6].

**Figure 3:**
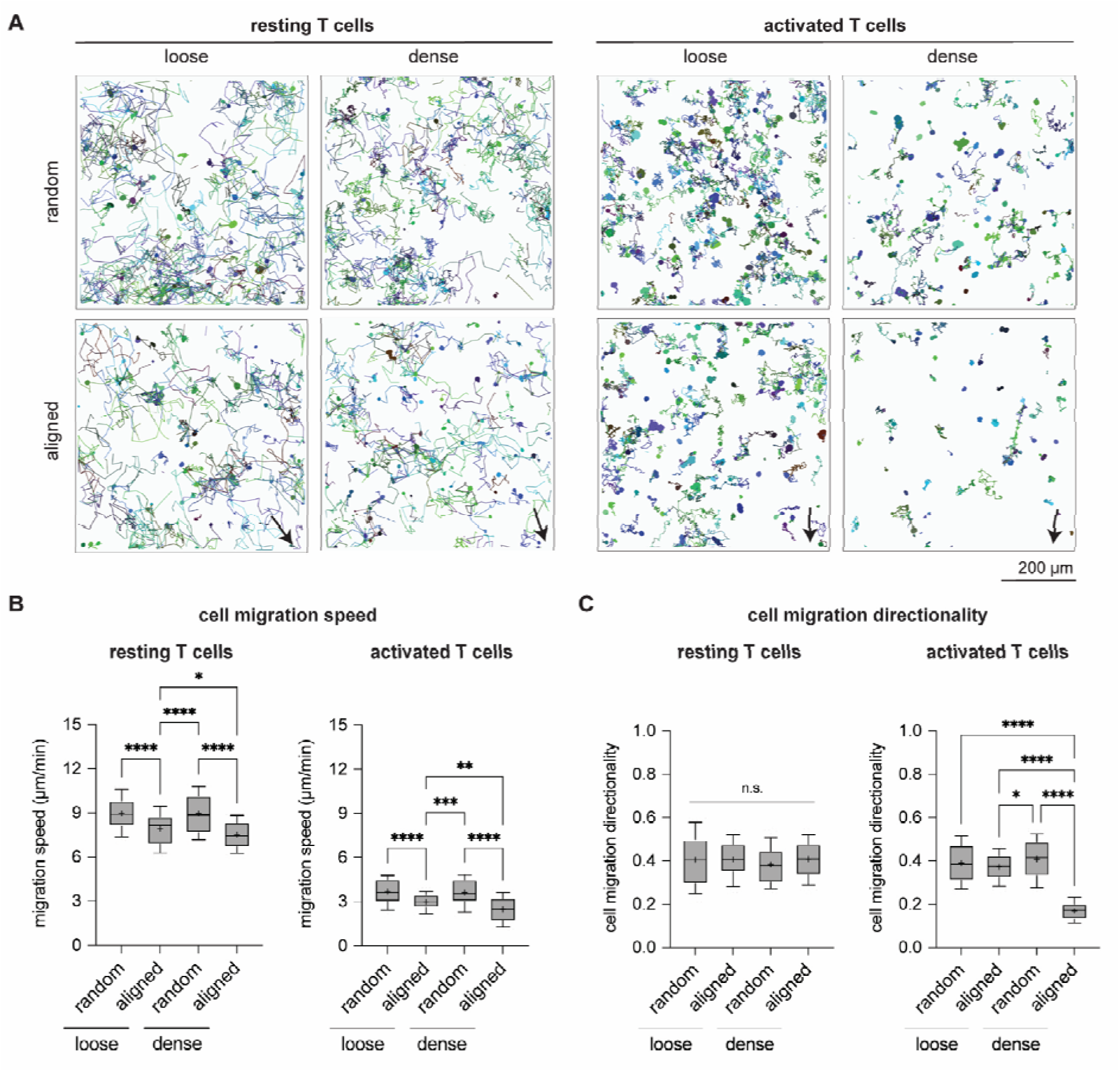
Migratory characteristics of resting and activated T-cells under different matrix conditions. Cell migration was observed at 6-minute intervals for 24 hours using a microscope equipped with a z-motor stage in bright-field mode. Cell migration trajectories were analyzed using stacked images with custom-built quantitative label-free 3D single-cell tracking software [35]. **(A)** Representative trajectories of resting and activated T-cells under different matrix conditions are depicted over 6 hours (scale bar = 200 µm). Each colored object and line represents a single cell and its migration trajectory. Cell migration characteristics were quantified in terms of **(B)** cell migration speed and **(C)** directionality for both resting and activated T-cells. At least 75 cells for each condition were analyzed. The data are presented as box plots with whiskers at the 10th and 90th percentiles. The ’+’ symbol and the horizontal line in the box plot indicate the mean and median of the data, respectively. Statistical significance was determined using one-way ANOVA with Dunn’s post hoc test, denoted as follows: *p < 0.05, **p < 0.01, ***p < 0.001, ****p < 0.0001. The experiments were conducted with four independent replicates.

To quantitatively confirm the observed differences in T-cell migration, we quantified the cell migration speed of both resting and activated T-cells under different matrix conditions, as depicted in **Figure 3B**. The migration speeds of resting T-cells in loose and dense random matrices were 8.9 ± 1.1 µm/min and 8.9 ± 1.3 µm/min, respectively. This finding suggested that the matrix density within the utilized pore size range of 6-11 µm did not significantly impact the amoeboid migration speed of resting T-cells. Interestingly, compared to those in random matrix conditions, resting T-cells in loose and dense matrices exhibited significant decreases in speed to 7.9 ± 1.1 µm/min and 7.5 ± 0.9 µm/min, respectively. On the other hand, activated T-cells exhibited reduced migration speeds in loose and dense random matrices, at 3.7 ± 0.9 µm/min and 3.6 ± 0.9 µm/min, respectively. Once again, the matrix density seemed to minimally affect the migration speed. However, in the aligned matrices, activated T-cells showed a significantly lower migration speed than in the random matrices, with speeds of 2.9 ± 0.5 µm/min and 2.4 ± 0.9 µm/min for the loosely and densely aligned matrices, respectively. Our findings suggest that despite similar pore sizes in both random and aligned matrices, there was a noticeable reduction in migration speeds for both resting and activated T-cells in aligned matrices, emphasizing the influence of matrix fibril orientation on T-cell migration.

In addition to cell migration, we further quantified the directionality of cell migration, as depicted in **Figure 3C**. Cell migration directionality quantifies the trajectory straightness by calculating displacement divided by the total path length, where a directionality score of 0 indicates random movement and a score of 1 signifies direct, straight-line migration from start to end-point. We observed no significant changes in the directionality of resting T-cells across all conditions. However, the directionality of activated T-cells was notably influenced solely by densely aligned tissue. Notably, studies tracking T-cell migration *in vivo* have highlighted how aligned fibers shape the migratory paths of T-cells within tumor microenvironments, guiding them along collagen fibers and leading to accumulation in the stroma [18, 53, 54]. Our findings corroborated well with a study using reconstituted collagen matrices with random, partially aligned, and aligned fibrils, which showed that collagen fibril alignment did not influence the directionality of T-cell migration [26]. However, another study reported that T-cell migration directionality is increased in aligned collagen matrices [13]. The discrepancy between these findings could be attributed to inadequate characterization of the matrix properties in the studies, which precludes direct comparison across different investigations. In addition, it should be noted that in both studies, cell tracking was conducted over a relatively short time period (20-30 minutes).

In summary, our findings indicate that the migration speeds of both resting and activated T-cell are influenced by collagen fibril alignment, while matrix porosity within the 6–11 µm range appears to exert a negligible effect on these speeds. We observed that T-cell migration directionality was significantly influenced within dense, aligned matrices but remained unaffected under the other matrix conditions tested in this study. Our results contribute to understanding the limited infiltration of T-cells into the tumor stroma, which is characterized by an alignment in the microarchitecture and denser tissue properties [18, 53].

### 3.4. Dense and aligned matrices reduced T-cell activation and proliferation

To determine whether alterations in the matrix microstructure affect T-cell activation, we quantified the early activation state of T-cells through the expression of cell surface markers and cytokines after 1 day of activation and subsequently investigated T-cell proliferation after 3 days of culture. **Figure 4A** shows the geometric mean fluorescence intensities (gMFIs) of CD25, CD44, CD69, and PD-1, which are major characteristics of early activated T-cells [3]. In general, the expression of these markers was significantly lower in resting T-cells than in activated T-cells (**Supplementary Figure S2**). The expression of these cell surface markers was significantly reduced in T-cells within dense random and aligned matrices (both loose and dense).

**Figure 4:**
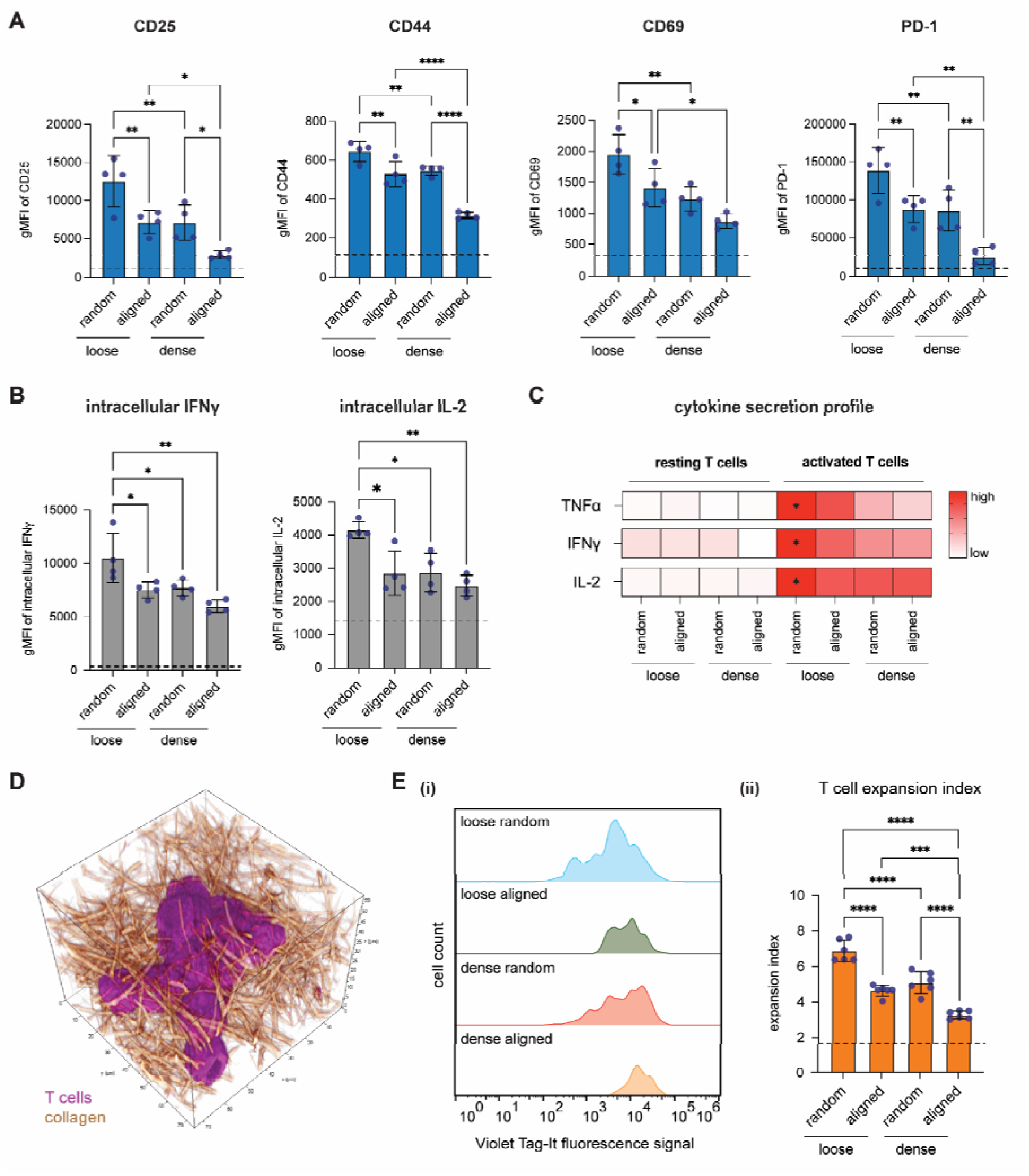
T-cell activation and proliferation under different matrix conditions. T-cell activation was investigated one day after activation by analysis. **(A)** The expression of cell surface markers, namely, CD25, CD44, CD69, and PD-1, was quantified by gMFI using flow cytometry. **(B)** The production of intracellular IFNγ and IL-2 was quantified via flow cytometry using the gMFI. For subfigures A and B, the dashed line represents the average gMFI for resting cells across all conditions. **(C)** A heatmap illustrating the quantification of secreted TNFα, IFNγ, and IL-2 using bead-based ELISA (red – high secretion, white - low secretion). * denotes comparisons where p < 0.05, indicating significant differences relative to all matrix conditions within groups. To assess T-cell proliferation, activated T-cells were cultured for three days post-activation. **(D)** A representative 3D image showing a T-cell cluster within a loose random matrix, with T-cells colored violet and collagen colored golden beige. **(E(i))** A representative histogram shows the Violet Tag-It fluorescence signal intensity of activated cells, analyzed using flow cytometry. **(E(ii))** Quantitative analysis of the EI via Violet Tag-It fluorescence intensity facilitates tracking of cell generations using FlowJo software’s proliferation module. The dashed line indicates the EI for resting cells across all conditions. The data are presented as the means ± SDs. Statistical significance was determined using one-way ANOVA with Dunn’s post hoc test; *p < 0.05, **p < 0.01, ***p < 0.001, ****p < 0.0001. The experiments were conducted with four independent replicates.

Additionally, upon T-cell activation, activated T-cells are required to produce three key cytokines: tumor necrosis factor alpha (TNFα), interferon gamma (IFNγ), and interleukin-2 (IL-2) [55, 56]. The production of TNFα induces apoptosis in infected cells and promotes leukocyte infiltration by regulating chemokine release and vascular permeability [57, 58]. IFNγ blocks microbial replication and enhances innate immune responses by directly activating macrophages [59] and recruiting neutrophils [60], while IL-2 promotes T-cell survival and proliferation [61]. We quantified the intracellular production of IFNγ and IL-2. As shown in **Figure 4B**, the production of the intracellular cytokines IFNγ and IL-2 was significantly reduced in activated T-cells within aligned (both loose and dense) and dense random matrices, demonstrating a trend similar to that of the expression of cell surface markers. We also quantified the intracellular cytokines production for resting cells, which was significantly lower than activated cells and we found no significant changes across all conditions (**Supplementary Figure S3**). Next, we quantified the secretion of TNFα, IFNγ, and IL-2 in the cell culture supernatant using bead-based ELISA. As shown in **Figure 4C**, we observed that TNFα and IFNγ, but not IL-2, were significantly reduced in activated T-cells within aligned (both loose and dense) and dense random matrices. Our data demonstrated that early activation levels were reduced in T-cells within dense random and aligned matrices (both loose and dense).

In addition, T-cell proliferation is a key parameter for measuring T-cell activation. **Figure 4D** shows a representative image of the T-cell cluster formed in collagen after 3 days of activation. To qualitatively investigate the proliferation of activated T-cells, we stained the cells with Violet-Tag-It dye, enabling tracking of T-cell generations through dye dilution, and monitored their proliferation over 3 days. We calculated the EI from the fluorescence signal intensity of the Violet-Tag-It dye, which represents the fold expansion of the cells. **Figure 4E(i)** shows the representative distribution of Violet-Tag-It fluorescence signal intensity in activated T-cells under different matrix conditions. Given that the Jurkat T-cells used were inherently proliferative, we first analyzed the EI of resting, nonactivated T-cells and observed that it remained constant across all matrix conditions (**Supplementary Figure S4**). The EI significantly decreased for activated T-cells within dense random and aligned matrices (both loose and dense), as demonstrated in **Figure 4E(ii)**. This decrease in the proliferation of activated T-cells correlated well with the previously discussed activation states of T-cells (**Figure 4A-C**). Our finding is in agreement with another study that demonstrated that increased collagen density can diminish T-cell proliferation [11, 62].Conversely, a study using 3D alginate hydrogels revealed enhanced T-cell proliferation in stiffer hydrogels [22]. However, alginate hydrogels lack the fibrillar microstructure of collagen matrices, which might be significant for translational purposes but limits their relevance for physiological or pathological models.

Overall, our data demonstrated suppressed T-cell activation and proliferation in dense random and aligned matrices (both loose and dense). This finding potentially reflects the immunosuppressive influence of the tumor microenvironment (TME) on T-cells and other immune cells.

### 3.5. Inhibition of YAP signaling improves T-cell activation

As the activation of T-cells within dense random and aligned matrices (both loose and dense matrices) was attenuated, we further hypothesized that mechanotransduction pathways might be triggered in resting T-cells, causing this suppressed activation. Therefore, we inhibited six mechanosensing and mechanotransduction biological pathways in resting T-cells prior to their activation: yes-associated protein (YAP), cell contractility, cytoskeleton, phosphoinositide 3-kinase (PI3K), piezo1 channel, and integrin interactions. We anticipated that blocking these pathways would restore T-cell activation within dense random and aligned matrices (both loose and dense matrices). To test this hypothesis, loosely aligned and dense random matrices were chosen. We quantified the expression of cell surface markers (CD25, CD44, CD69, and PD-1) in activated T-cells using flow cytometry. As shown in **Figure 5A**, only YAP inhibition enhanced the activation of T-cells, while other inhibitors had little or no significant effect on activation. This result indicates that YAP activity is increased in resting T-cells, which in turn results in the suppression of T-cell activation.

**Figure 5:**
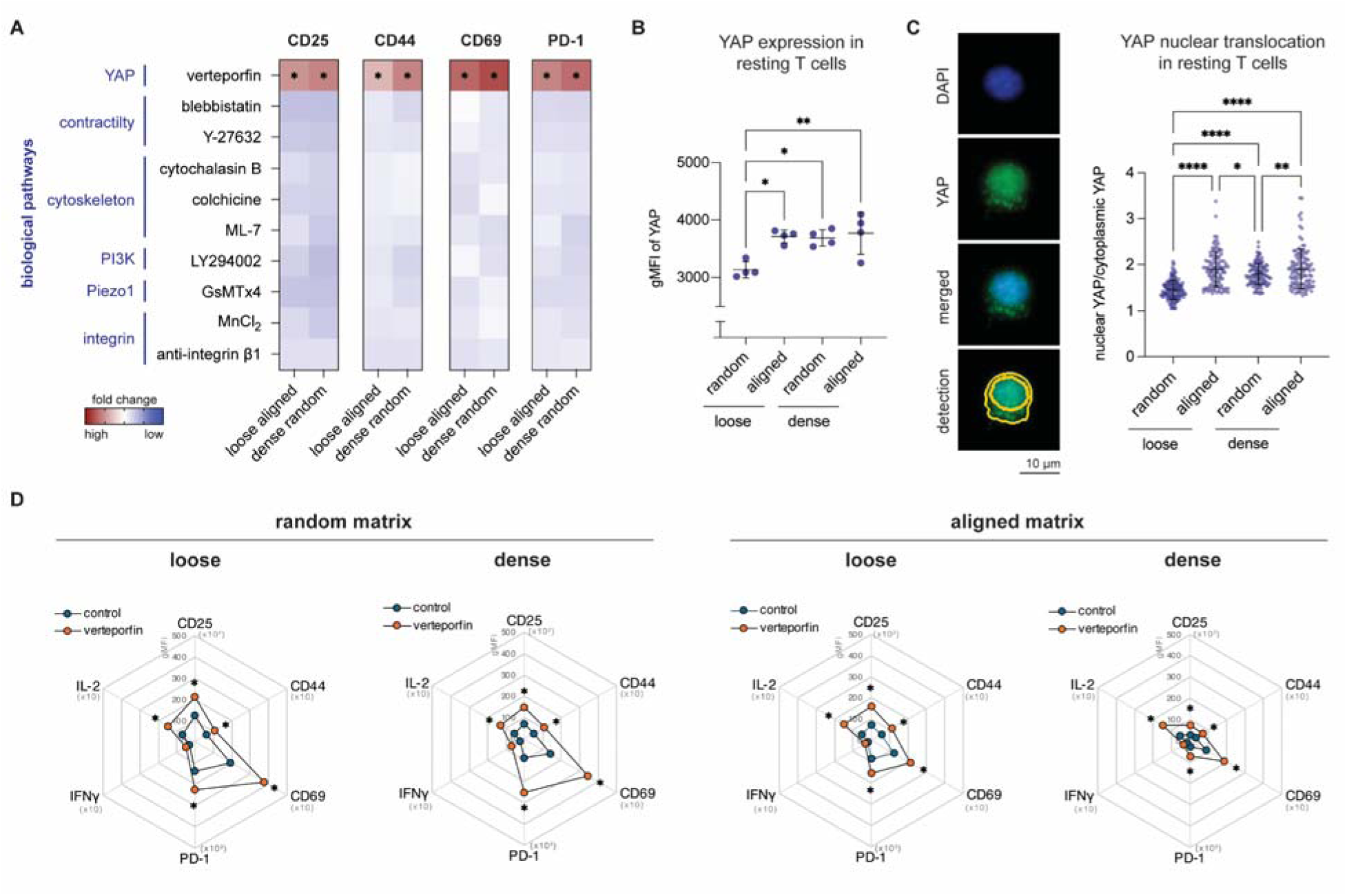
Inhibition of mechanotransduction pathways associated with the suppression of T-cell activation. Prior to T-cell activation, resting T-cells were treated with various inhibitors for 3 hours. Subsequently, the cells were chemically activated using PMA and ionomycin for 3 hours followed b fresh media for 24 hours. **(A)** A heatmap illustrating the fold change in the expression of treated T-cell activation markers under loosely aligned and dense random matrix conditions normalized to its non-treated counterpart, quantified using flow cytometry (red indicates high expression, blue indicates low expression). **(B)** Quantitative analysis of the gMFI of YAP expression in resting cells was performed using flow cytometry. **(C)** YAP nuclear translocation in resting T-cells was analyzed using the built-in analysis software BioTek Lionheart FX. The analysis segmented the cytoplasm and nucleus based on the YAP fluorescence signal and the DAPI signal, respectively. The fluorescence intensity of YAP in the segmented cytoplasm and nucleus was calculated as a ratio. At least 109 cells for each condition were analyzed. For subfigures B and C, the data are presented as the means ± SDs. The dashed line in subfigure C represents the average gMFI for resting cells across all conditions. **(D)** Quantitative analysis of T-cell activation markers and intracellular cytokine production upon YAP inhibition prior to activation under all matrix conditions. The means of the data are presented. Statistical significance was determined using one-way ANOVA with Dunn’s post hoc test; *p < 0.05, **p < 0.01, ***p < 0.001, ****p < 0.0001. The experiments were conducted with four independent replicates.

To confirm that YAP signaling is triggered in resting T-cells, we investigated YAP expression and translocation. As shown in **Figure 5B**, increased expression of YAP was observed in resting T-cells within dense random and aligned matrices (both loose and dense matrices). It is well known that phosphorylated YAP remains within the cytoplasm, and once signaling is activated, YAP is dephosphorylated and translocated into the nucleus, where it interacts with transcriptional enhancer associate domain (TEAD) family proteins [63, 64]. We therefore analyzed YAP translocation in resting T-cells under different matrix conditions. Cytoplasmic and nuclear YAP (areas overlapping with DAPI) were quantified, as shown in representative images in **Figure 5C**. The results suggested that the ratio of nuclear to cytoplasmic YAP increased in resting T-cells within dense random and aligned matrices (both loose and dense matrices). These findings indicate a greater translocation rate to the nucleus and, consequently, greater YAP activity in these matrices. These results are corroborated by previous studies showing that YAP knockout in mouse T-cells increases their activation state and improves their function in tumor microenvironments [65–67].

To assess whether YAP inhibitors boost T-cell activation, we inhibited YAP signaling across diverse matrix conditions and evaluated its activation by analyzing the expression of cell surface markers and the production of intracellular IFNγ and IL-2. **Figure 5D** shows that YAP inhibition markedly elevated the expression levels of CD25, CD44, CD69, and PD-1, as well as the production of intracellular IL-2, across all tested conditions. Conversely, the influence on intracellular IFNγ production was minimal. Notably, YAP inhibition increased T-cell activation in dense, aligned matrices to levels similar to those observed in loose, random matrices, with significantly increased intracellular IL-2 production (**Supplementary Figure S5**). Moreover, the effect of YAP inhibition was more significant in T-cells within random matrices than in their aligned counterparts. These findings suggest the possible role of an additional mechanotransduction pathway that, along with YAP, might suppress T-cell activation, warranting further investigation.

In our study, we showed that YAP signaling is a key player in suppressing T-cell activation within dense and aligned matrices. However, we were unable to pinpoint the exact external signals that activate YAP. Previous research suggested that YAP can be activated by mechanisms such as actomyosin contractility [68, 69], Piezo1 channels [70–72], and β1 integrin interactions [73–75]. Despite thes reported insights, our experiments indicate that blocking these potential activators does not restore T-cell activation (**Figure 5A**). Future investigations should aim to uncover the specific external signaling pathways that influence YAP activity in T-cells. Such discoveries could greatly enhance the effectiveness of adoptive T-cell immunotherapy by providing new therapeutic targets.

## 4. Conclusion

In this work, we investigated how the biophysical characteristics of the ECM influence T-cell behavior, focusing on the impact of ECM density and alignment on T-cell activation and function. Utilizing biomimetic 3D collagen matrices with controlled alignment and density, we demonstrated that T-cells exhibited decreased activation, reduced cytokine production, and limited proliferation within dense and aligned ECM matrices. These observations underscore the critical role of physical cues from the ECM in modulating T-cell responses, thereby enhancing our understanding of T-cell mechanobiology. Furthermore, our study revealed that YAP signaling is a key mechanotransduction pathway involved in suppressing T-cell activation. The inhibition of YAP signaling enhances T-cell activation. These results indicate a connection between the biophysical properties of the ECM and mechanosensing through YAP in T-cells. This insight not only advances our fundamental understanding of T-cell biology but also highlights a potential avenue for therapeutic intervention, particularly in adoptive T-cell immunotherapy.

## Acknowledgments

The authors acknowledge support from the New York University Abu Dhabi (NYUAD) Faculty Research Fund (AD266) and the New York University Abu Dhabi Global Ph.D. Fellowship. The authors would also like to acknowledge support from the NYUAD core technology platform: Light Microscopy and Molecular and Cell Biology.

## Data availability

The datasets generated during and/or analyzed during the current study are available from the corresponding author upon reasonable request.

## Code availability

A custom-built MATLAB script for measuring collagen microarchitecture can be found at https://git.sc.uni-leipzig.de/pe695hoje/topology-analysis.

