## Supplemental Information for "Matrix alignment and density modulate YAP-mediated T-cell immune suppression"

Prof. Dr. Jeremy Teo

| | |
|---|---|
| Postal address | New York University Abu Dhabi<br>Division of Engineering<br>Abu Dhabi, UAE. |
| Telephone | +971 2 6286689 |
| E-mail: | |

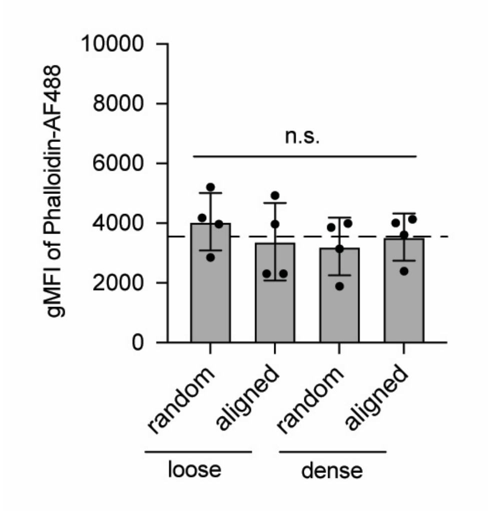


**Figure S1: Actin content in resting T-cells under different matrix conditions.** Quantitative analysis of actin content in resting T-cells was performed using flow cytometry to measure the gMFI of phalloidin-Alexa Fluor 488-stained cells. The data are presented as the means ± SDs. Statistical significance was assessed using one-way ANOVA with Dunn's post hoc test; no significant differences were detected across the matrix conditions. The experiments were conducted with four independent replicates.

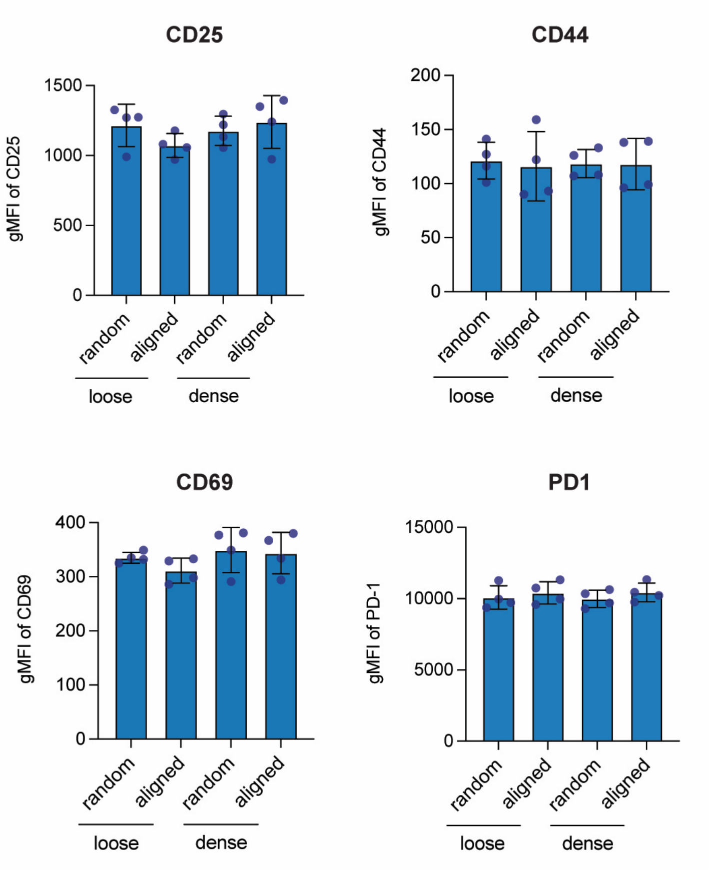


**Figure S2: Expression of T-cell activation markers in resting T-cells within different matrix conditions.** The expression of cell surface markers, namely, CD25, CD44, CD69, and PD-1, was quantified by gMFI using flow cytometry. The data are presented as the means ± SDs. Statistical significance was assessed using one-way ANOVA with Dunn's post hoc test; no significant differences were detected across the matrix conditions. The experiments were conducted with four independent replicates.

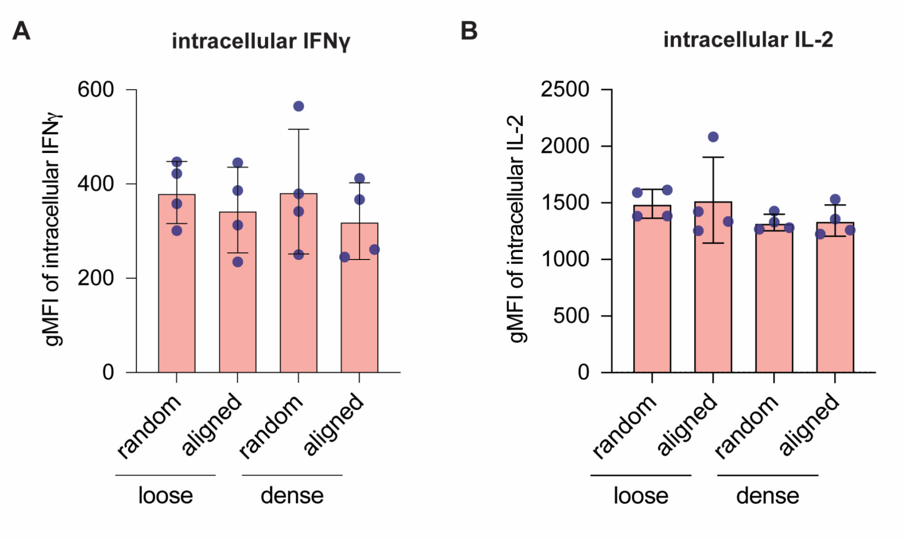


**Figure S3: Intracellular IFNγ and IL-2 production in resting T-cells under different matrix conditions.** The production of intracellular IFNγ and IL-2 in resting T-cells was analyzed through intracellular staining and quantified using gMFI via flow cytometry. The data are presented as the means ± SDs. Statistical significance was assessed using one-way ANOVA with Dunn's post hoc test; no significant differences were detected across the matrix conditions. The experiments were conducted with four independent replicates.

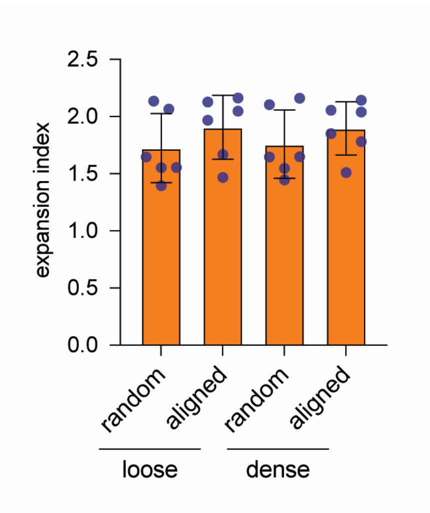


**Figure S4: Expansion indices of resting T-cells under different matrix conditions.** Quantitative analysis of the EI of resting T-cells, determined by measuring Violet-Tag-It fluorescence intensity, was performed using FlowJo software's proliferation module. The dashed line represents the EI for resting cells across all conditions. The data are presented as the means ± SDs. Statistical significance was assessed using one-way ANOVA with Dunn's post hoc test; no significant differences were found across the matrix conditions. The experiments were conducted with four independent replicates.

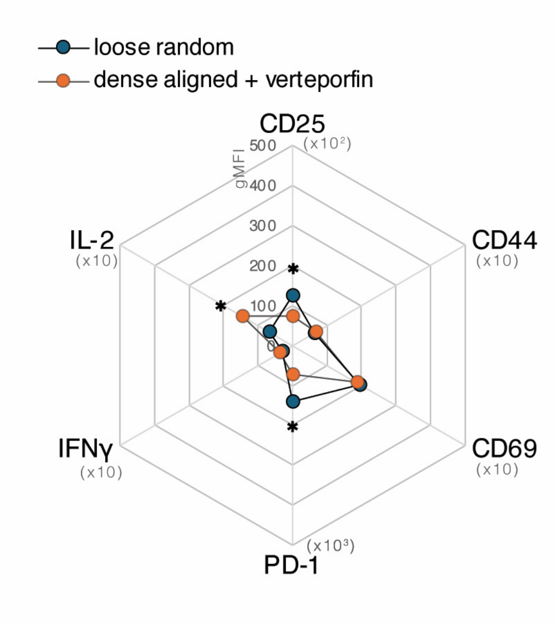


**Figure S5: Inhibition of YAP enhances T-cell activation in a densely aligned matrix.** Figure 5D shows that YAP inhibition enhances T-cell activation in a dense aligned matrix, achieving levels comparable to those observed in a loose random matrix (without YAP inhibition). Quantitative analysis of T-cell activation markers and intracellular cytokine production following YAP inhibition prior to T-cell activation is shown. The data are presented as the means. Statistical significance was assessed using one-way ANOVA with Dunn's post hoc test, indicated by *$p < 0.05$. The experiments were conducted with four independent replicates.
